# The NFκB *Dif* is required for behavioral and molecular correlates of sleep homeostasis in *Drosophila*

**DOI:** 10.1101/2023.10.12.562029

**Authors:** Michael K. O’Hara, Christopher Saul, Arun Handa, Amita Sehgal, Julie A. Williams

**Affiliations:** Chronobiology and Sleep Institute, Department of Neuroscience; Howard Hughes Medical Institute, University of Pennsylvania Perelman School of Medicine Philadelphia, PA 19104

**Keywords:** Dif, Relish, NFκB, sleep deprivation, *nemuri*

## Abstract

The nuclear factor binding the κ light chain in B-cells (NFκB) is involved in a wide range of cellular processes including development, growth, innate immunity, and sleep. However, efforts have been limited toward understanding how specific NFκB transcription factors function in sleep. *Drosophila* fruit flies carry three genes encoding NFκB transcription factors, *Dorsal*, *Dorsal Immunity Factor* (*Dif*), and *Relish*. We previously found that loss of the *Relish* gene from fat body suppressed daily nighttime sleep, and abolished infection-induced sleep. Here we show that *Dif* regulates daily sleep and recovery sleep following prolonged wakefulness. Mutants of *Dif* showed reduced daily sleep and suppressed recovery in response to sleep deprivation. Pan-neuronal knockdown of *Dif* strongly suppressed daily sleep, indicating that in contrast to *Relish*, *Dif* functions from the central nervous system to regulate sleep.

Based on the distribution of a *Dif*-associated GAL4 driver, we hypothesized that its effects on sleep were mediated by the pars intercerebralis (PI). While RNAi knock-down of *Dif* in the PI reduced daily sleep, it had no effect on the recovery response to sleep deprivation. However, recovery sleep was suppressed when RNAi knock-down of *Dif* was distributed across a wider range of neurons. Induction of the *nemuri* (*nur*) antimicrobial peptide by sleep deprivation was suppressed in *Dif* mutants and pan-neuronal over-expression of *nur* also suppressed the *Dif* mutant phenotype. Together, these findings indicate that *Dif* functions from brain to target *nemuri* and to promote sleep.

**Statement of Significance:** NFκB transcription factors drive inflammatory processes that underly a multitude of human diseases that are accompanied by sleep disturbance. However, genetic studies in mammals that investigate a function of NFκB in sleep have been limited. Using genetic approaches in the *Drosophila* fruit fly, we show that the *Dif* NFκB, homologous to the *Rel* family of NFκBs in humans, functions from neuronal tissue to regulate daily sleep and to mediate responses to sleep loss. These findings enhance our understanding of the role of a specific NFκB gene in sleep regulation.

## Introduction

A connection between sleep and the immune system has long been implicated ^1–3^, but the nature of this connection is not well understood. Despite its central role in inflammation and illness ^4,5^, very few genetic studies in rodent models have investigated a function of specific NFκB genes in daily sleep ^6,7^. To address this gap, we have previously explored the role of NFκB in sleep using the *Drosophila* genetic model ^8,9^, where NFκB is also central to innate immune function ^5,10,11^.

Both mammalian and *Drosophila* species carry multiple genes encoding NFκB transcription factors from one of two families ^4,12^. One gene family consists of NFκB1 (p105) and NFκB2 (p100), which are processed into the active forms, p52 and p50, respectively, in response to stress or other stimuli. These two genes are homologous to *Relish* in *Drosophila*. Jhaveri and colleagues ^6^ reported that p50 knockout mice showed increased daily sleep, but were defective in initiating sleep induced by immune challenge, particularly in response to the influenza virus. In contrast to the mouse model, *Relish* mutants reduced daily nighttime sleep in heterozygous females ^8^. However, like the mouse model, sleep induced by either septic or aseptic injury was abolished in homozygous null mutants of *Relish* ^9^. These findings in mammals and flies indicate a conserved function of the NFκB1/2 gene family in infection-induced sleep.

The second NFκB gene family, “Rel”, consists of three genes in mammals, RelA, RelB, and c-Rel^4^. These encode proteins that interact with an inhibitory IκB and become active when this regulatory protein is targeted for degradation. Dorsal (*dl*) and *dorsal immunity factor* (*Dif*) are the *Drosophila* orthologs for this family and are targeted by the Toll receptor signaling cassette, which includes the IκB Cactus ^13,14^. Dorsal was initially discovered as a factor that controls embryonic patterning during development ^15^, and its role in adult immune function is unknown. In contrast to its paralog, the function of *Dif* in the innate immune response is well established. While targeted by the same signaling pathways, *Dif* mediates larval and adult immune responses to gram positive bacteria and fungi ^11^.

Here we investigate a role of *Dif* in daily sleep. In contrast to *Relish*, *Dif* mutants have reduced daily sleep in both males and females. Flies showed defects in response to sleep deprivation such that there were long delays to initiating recovery sleep, which itself was reduced as compared to wild type controls. While *Relish* regulates sleep from fat body ^8,9^, in the current study we find that *Dif* regulates sleep from multiple cell types, including fat body and neurons. However, its role in recovery sleep after sleep loss is neuronally mediated. *Dif* is known to target numerous antimicrobial peptides ^16,17^. We therefore evaluated its function in activating expression of the *nemuri* antimicrobial peptide, which promotes sleep ^18^. *Nemuri* is dependent on DIF for its induction with sleep deprivation, and overexpression of *nemuri* suppressed the effects of *Dif* on daily sleep. These findings indicate distinct roles of NFκB in regulation of sleep and identify a mechanism by which *Dif* promotes sleep.

## Methods

### Fly stocks

Male and female *Drosophila melanogaster* were used for the studies as indicated. *Dif^1^*, *Relish^E20^*, and *Dif^1^*; *Relish^E20^*mutants were the same as those used in Kuo et al (2014). Briefly, flies were backcrossed to a Canton-S wild type strain for eight generations and compared to the original Canton-Special wild type as control. *UAS-nemuri* was as described previously ^18^. *Dif* RNAi lines were obtained from the Vienna Drosophila Research Center, #30579 and #100537. Gal4 lines were also obtained from the Bloomington *Drosophila* Stock Center (BDSC) or were available from the lab stocks: *elav-geneswitch* ^19^, *actin-Gal4* (BDSC #3954), *Dif-Gal4* (BDSC #64720), *Dilp2-Gal4*, *c767-Gal4*, *Gbp3-Gal4* (BDSC #64111), *Repo-Gal4*, *c309-Gal4*, *201y-Gal4*, *ppk-Gal4*, *hugin-Gal4*, Vienna Tile Gal4 lines ^20^: VT050161, VT018146, VT014615, VT003282, VT036268, VT008975, and VT056377, and *UAS-GFP:Cd8* ^21^. For analysis of RNAi, where applicable, the Gal4 line was crossed to wild type *w^1118^* flies (the background for the *UAS-RNAi-Dif* flies) as one control, and the wild type background of the Gal4 line (either *w^1118^* or *Iso^31^*) was crossed to *UAS-RNAi-Dif* as the other. Experimental test lines were offspring from the *Gal4* line crossed with *UAS-RNAi-Dif*.

### Behavioral Assays

All flies were reared on a standard corn-meal medium as described on the BDSC website (soy flour and light corn-syrup-based). Flies were maintained in the same light: dark schedule used for behavioral assays. Locomotor activity and sleep were measured as described previously ^22^. Briefly, male and female flies that were 3-5 days of age were loaded into glass activity tubes (65 mm in length) containing 5% sucrose, 2% agar medium at one end and plugged with cotton/acrylic yarn at the other end. Glass tubes were loaded into Drosophila Activity Monitors (DAM2, multibeam MB5 or DAM5H; Trikinetics, Inc., Waltham, MA), and activity was recorded for 5-7 days. Monitors were placed in incubators kept at 25° C in a 12h:12h light: dark cycle and 50% humidity. In this assay, activity counts correspond to breaks of one or more IR beams that bisects the tube. Sleep is defined as an activity count of zero for a minimum of 5 consecutive minutes ^23^. Sleep latency was determined from the time of lights-on (for recovery sleep) and from the time of lights-off (for nighttime sleep). Flies that were already asleep from before these time points were excluded from the analysis of latency values (in minutes). Data were processed using custom software, Insomniac 3 ^24^. For each locomotor assay, 8 flies (for MB5 monitors) to 16 flies (for DAM5H or DAM2 monitors) per group were loaded into monitors for each of 2-4 experimental replicates.

For drug dependent induction of *Dif*-RNAi, 25 μM mifepristone (RU486, Sigma-Aldrich, St. Louis, MO) was added to standard fly food medium for chronic exposure throughout development^19,25^. RU486 was diluted from a 10 mM stock in 80% ethanol. Equivalent dilutions were made with 80% ethanol for the vehicle control groups. Adult flies were exposed to 500 μM RU486, or equivalent control vehicle dilution, in sucrose/agar medium used in the activity tubes for behavioral experiments. For drug dependent induction of *UAS-nemuri*, adult flies were fed 500 μM RU486 on sucrose-agar medium as reported previously^18^.

To sleep deprive flies, up to four DAM2 monitors were attached to a multi-tube vortexer (VWR) fitted with a mounting plate (Trikinetics, Inc.). The vortexer was attached to an LC4 light controller and programmed to apply a 1 second pulse at random intervals from 2-40 seconds for up to 8 hours.

### Immunohistochemistry

Female flies carrying *Dif-Gal4>UAS-GFP:CD8* were subjected to 6h sleep deprivation or left undisturbed and collected at ZT 0 (ZT = zeitgeber time). Whole brains were dissected out in cold phosphate buffered saline, fixed in 4% paraformaldehyde, rinsed and processed with a rabbit anti-GFP antibody (1:1000, Life Tech #A-11122). Secondary labeling was with anti-rabbit Alexa488 (1:1,000, ThermoFisher# A-11008). Brains were then mounted with Vectashield medium (Vector Laboratories, H-1200-10) and visualized using a laser-scanning confocal microscope (Leica SP5).

### Q-PCR

To evaluate expression of *Dif* mRNA in RNAi lines, flies of both sexes of indicated genotypes (Fig S4) were collected on dry ice at the time of lights-on, zeitgeber time (ZT) 0. Total RNA was extracted from whole flies. To evaluate expression of *nemuri* mRNA, female flies of indicated genotypes (Fig 7A) were sleep deprived for 6h as described above and were harvested on dry ice at ZT 0 along with undisturbed controls. Heads were removed and collected on dry ice. At least three biological replicates were collected for each genotype and treatment groups.

Total RNA was extracted from all samples using Trizol reagent (Ambion; Sigma Aldrich). Residual genomic DNA was removed from samples containing 2 μg of total RNA using the Turbo DNase-free kit (Invitrogen; Thermofisher Scientific) and about 1.8 μg of total RNA was reverse transcribed using a High Capacity RNA to cDNA kit (Applied Biosystems; Thermofisher Scientific). Additional samples in which RNA was not reverse transcribed were included as controls and indicated effective genomic DNA removal.

Amplification and detection of *Dif*, *nemuri* and a reference gene, *tubulin*, were accomplished using the PowerUp SYBR Green Master Mix by Applied Biosystems and target-specific primers. Primers used to amplify *nemuri* and *tubulin* were previously described by Toda et al. ^18^ and Xu et al. ^26^, respectively (*nemuri*: forward 5’ ACGTCGGACAACAAGGAAGACTC 3’, reverse 5’ CCTCGGCAACATCATCGCTG 3’; *tubulin*: forward 5’- CGTCTGGACCACAAGTTCGA 3’, reverse 5’ CCTCCATACCCTCACCAACGT 3’). Primers used to amplify *Dif* were forward: 5’ CGTTCGAGTACTACCCGAATCC-3’, reverse: 5’ CTTTAGGCTTTCAACTGTTTTTTGG- 3’. Reactions were carried out in the ViiA7 Real-Time PCR System (Applied Biosystems) with an initial 10-minute denaturation step at 95°C followed by 40 cycles of a 15-second denaturation step at 95°C and a 1-minute annealing and extension step at 60°C.

To assess the capacity for *Actin-Gal4*-directed *Dif^RNAi^ ^#30579^* and *Dif^RNAi^ ^#100537^* expression to reduce *Dif mRNA* expression, primer pair-specific efficiencies were used to weight qPCR cycle threshold (Ct) values before normalizing to *tubulin*, as described by Ganger *et al*. (2017)^27^. Normalized ΔΔCt values were used for statistical analyses and to estimate *Dif mRNA* quantity. *Dif^RNAi^ ^#30579^* and *Dif^RNAi^ ^#100537^* were analyzed separately.

To assess the expression of *nemuri* mRNA in response to sleep deprivation, transcript quantity for each genotype was calculated relative to the indicated control group using the comparative C_T_ method (2^-ΔΔCT^; ^28^).

### Statistical Analyses

One-way ANOVA followed by Tukey’s post-hoc was used to compare overall sleep parameters in 12h or 24h increments, and for comparisons between groups for changes in mRNA expression of *Dif* or *nemuri* mRNA. Non-parametric tests were used for data that did not fit a normal distribution as determined by the Shapiro-Wilk test, particularly for sleep latency, and for normalized data (% gain, % sleep loss, and 24 hour sleep values). For experiments using RU486-dependent gene-switch lines, two-way ANOVA was used to evaluate interactions of drug and genotype dependent effects on sleep.

To quantify recovery sleep from deprivation (sleep gain), total daytime sleep (minutes/12 hours) before the night of deprivation (D_b_; “b” = baseline) was subtracted from total sleep during the daytime period after deprivation (D_r_; “r” = recovery). Likewise, sleep loss was determined by subtracting the baseline nighttime period (N_b_) from the 12h nighttime period that flies were subjected to deprivation (N_d_; “d”= deprivation). Percent sleep loss was thus calculated: │N_d_-N_b_│/N_b_. Percent sleep gain was calculated as described in Cirelli et al ^29^, where the minutes sleep gained (D_r_-D_b_) was divided by the absolute value of minutes of sleep lost (N_d_-N_b_). Percent sleep gain, percent sleep loss, and sleep latency data are depicted as box-and-whisker plots. Outliers were defined as values greater than 1.5 times the inter-quartile range. A Kruskal-Wallis test followed by Dunn’s post-hoc (with Bonferroni corrected p-values) was used to evaluate effects of genotype as indicated.

Flies of both sexes were used in most experiments but were analyzed separately due to the known differences in sleep behavior between male and female flies. Sleep deprivation experiments were in female flies only because males show limited responsiveness to sleep loss^23^.

All statistical analyses were performed using public-accessed software (PAST; http://folk.uio.no/ohammer/past/) ^30^.

## Results

### Mutations in *Dif* reduce sleep

We first determined daily sleep in *Dif^1^* mutant flies. The *Dif^1^* allele is a point mutation that results in a change of a glycine residue (181) to aspartic acid in part of the DNA binding domain. This change is predicted to hinder the ability of the DIF protein to bind DNA, thus rendering the mutant a hypomorph ^17^. Both male and female *Dif^1^* mutants showed reduced daily sleep, with robust effects at night (Fig 1A-D). We previously showed that null mutants of another NFκB, *Relish* (*Rel^E20^*)^31^ reduced nighttime sleep in female but not in male flies, and did not affect responses to sleep loss ^8,32^. To test whether loss of the *Relish* gene was additive to the effects of *Dif*, we compared *Dif^1^* mutants to those carrying both *Dif^1^* and *Rel^E20^* alleles. The double mutant further reduced sleep in female but not in male flies. The reduction of sleep in all flies was attributed to reduced nighttime sleep, with a significantly shortened sleep bout duration, suggesting either a reduced need for sleep or an inability to initiate and/or maintain sleep (Fig 1A-1F). In support of the former hypothesis, latency to the first nighttime sleep bout was significantly increased in *Dif^1^* and *Dif^1^*; *Rel^E20^*double mutants (Fig 1G).

**Figure 1.**
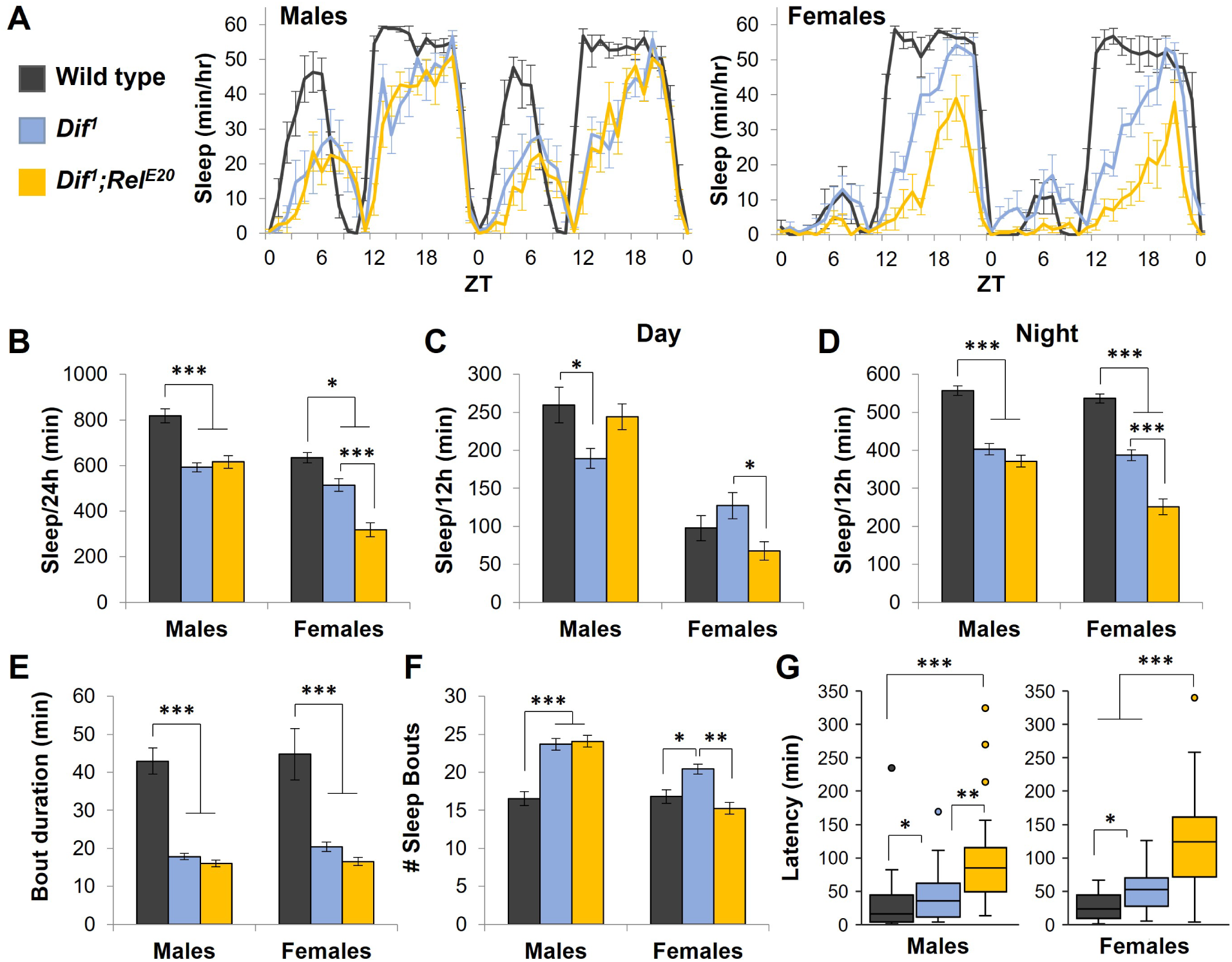
NFκB mutants reduce daily sleep. **A**. Representative experiment showing sleep (mean ± SEM; n=8 flies each group) per hour plotted for wild type, *Dif^1^* and *Dif^1^; Rel^E20^* double mutant flies. ZT = zeitgeber time, where ZT 0-12 corresponds to lights-on, and ZT 12-24 lights off. Legend key applies to A-G. Mean ± SEM daily sleep (24h; **B**), for daytime (**C**), and nighttime (**D**) is shown for male (n=38-40) and female (n=39-40 per group) flies. Mean ± SEM is shown for nighttime sleep bout duration (**E**) and number of bouts (**F**); ***= p<0.00001, **=p<0.001, * = p<0.05, ANOVA with Tukey’s post-hoc. **G**. Box and whisker plots with outliers showing latency to nighttime sleep onset in indicated groups. *** = p<0.00001, **=p<0.001, *=p<0.05, Kruskall-Wallis with Dunn’s post-hoc (Bonferroni corrected); n=37-40 flies per group.

To determine whether the reduced sleep phenotype mapped to the chromosomal locus that included the *Dif* gene, we crossed *Dif^1^* mutants to those that contained a GAL4 driver sequence inserted in to 5’ end of the *Dif* gene (*Dif^G4^*)^33^, or to an equivalent control background, *w^1118^*. Given the position of the *Gal4* sequence, we hypothesized that it would disrupt expression of the *Dif* gene. Total daily sleep was strongly reduced in the *Dif^1^*/*Dif^G4^* transheterozygotes indicating that the *Dif^1^* allele failed to complement *Dif^G4^*(Fig 2A, 2B). Heterozygous mutants of both alleles showed semi-dominant phenotypes such that total sleep was modestly but significantly reduced in males, whereas sleep in female flies was more sensitive to the *Dif^G4^/+* allele. Nighttime sleep architecture showed significant reduction in sleep bout length (Fig 2C) with intermediate reduction in the heterozygous flies. The number of sleep bouts was increased by all alleles in males but decreased in the transheterozygous females relative to wild type controls (Fig 2D). Finally, similar to the *Dif1* homozygous flies, transheterozygous *Dif^1^*/*Dif^G4^*mutants showed a significant delay to nighttime sleep onset (Fig 2E).

**Figure 2.**
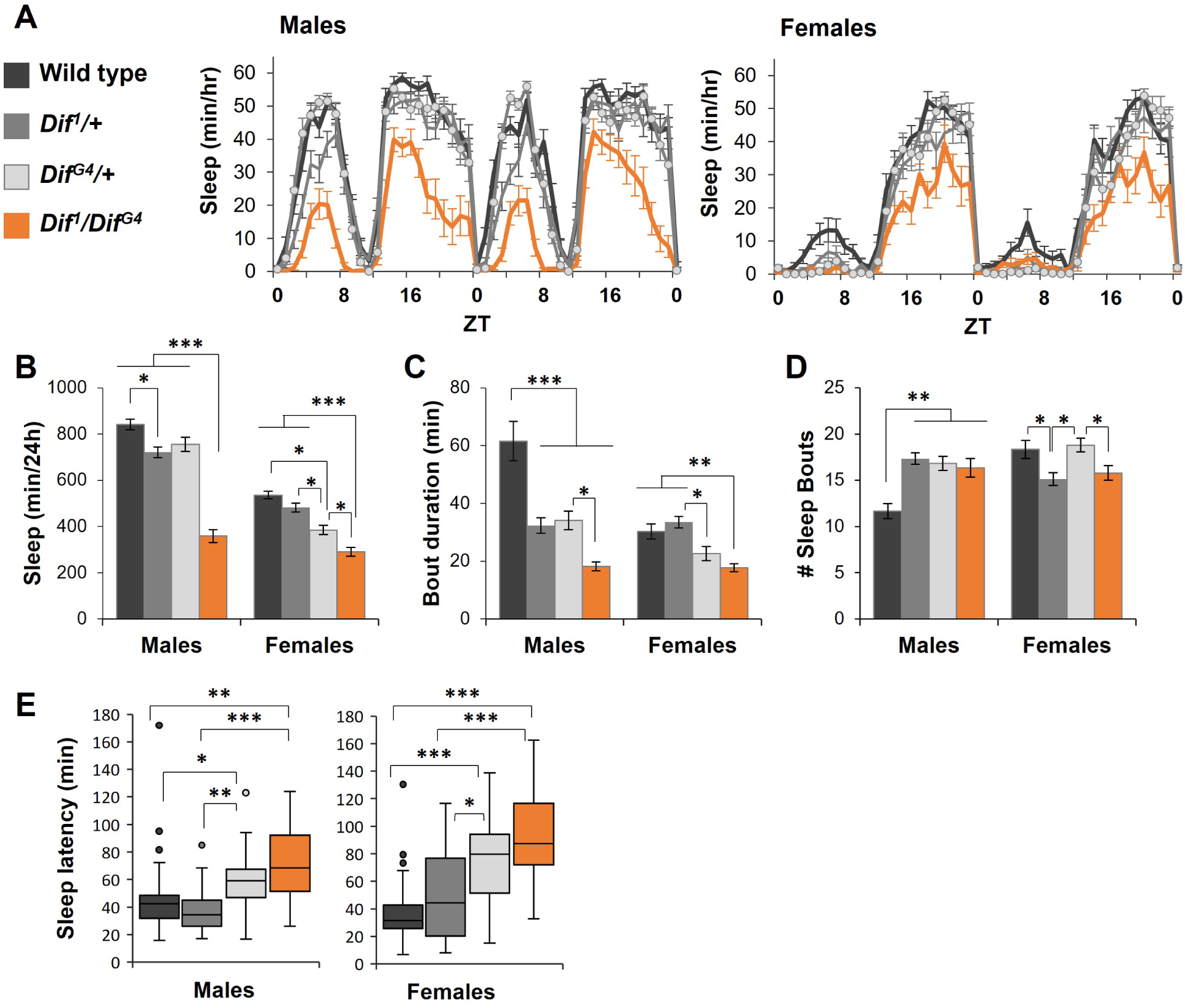
Transheterozygous mutants of *Dif* reduce daily sleep. **A**. Representative experiment showing sleep (mean ± SEM; n=16 flies each group; ZT = zeitgeber time) per hour plotted for wild type, *Dif^1^/+*, *Dif^G4^/+*, and *Dif^1^/Dif^G4^* flies. Legend key applies to all figure panels. Mean ± SEM daily sleep per 24h is shown in (**B**), nighttime sleep bout duration (**C**), and number of nighttime sleep bouts (**D**); ***= p<0.00001, **=p<0.001, * = p<0.05, ANOVA with Tukey’s post-hoc. **E**. Box and whisker plots showing latency to nighttime sleep onset. *** = p<0.00001, **=p<0.001, *=p<0.05, Kruskall-Wallis with Dunn’s post-hoc (Bonferroni corrected); n=32 flies per group.

We also evaluated whether the *Dif^1^* allele complemented a chromosomal lesion at the *Dif* locus, *Df(2L)J4* (J4; ^34^). *Dif^1^/J4* flies were compared to *J4/+* and CS wild type flies. The *Dif^1^* allele failed to complement the *J4* deficiency, as transheterozygous flies showed reduced daily sleep (Fig S1). Nighttime sleep architecture showed modest but significant effects relative to one or both control lines, with reduced sleep bout duration and increased time to sleep onset. Together these findings indicate that the DIF NFκB promotes daily sleep in both male and female flies.

### Recovery sleep is reduced in *Dif* mutants

To test whether *Dif* and/or *Relish* signals contribute to sleep need, we next subjected NFκB mutants, *Rel^E20^*, *Dif^1^*, and *Dif^1^; Rel^E20^* double mutants, to nighttime sleep deprivation. Flies were mechanically stimulated for 8h from ZT 16 until lights-on (Fig 3A). While *Rel^E20^*mutants showed the highest net sleep gain relative to other groups during the recovery period, the daytime sleep gain (as a function of baseline; see methods) was significantly reduced in both *Dif^1^* and *Dif^1^;Rel^E20^* double mutants (Fig 3B). However, a confounding factor is the fact that both mutants have less sleep at night, and so they may be deprived of less sleep, thereby requiring less recovery. To circumvent this issue, we first calculated the total nighttime sleep lost as a percentage of the previous undisturbed night. As the mutants experience most of their sleep during the latter half of the night, the median percent sleep loss was greater in flies lacking *Dif* as compared to CS wild type or *Rel^E20^*homozygous mutants (Fig 3C). In contrast, correcting for differences in baseline sleep, the percent sleep gain following deprivation in *Dif^1^* flies was significantly lower as compared to all other groups (Fig 3D). Moreover, the amount of time to onset (latency) of the first daytime sleep bout was also significantly increased (Fig 3E). In *Dif^1^/ Dif^G4^* transheterozygous mutants, we did not detect significant changes in % sleep gain after deprivation (Fig 3F). However, sleep onset during the recovery period was significantly delayed (Fig 3G). Similar results were found in *Dif^1^/J4* transheterozygous mutants, such that sleep gain following deprivation was reduced, and onset to recovery sleep was increased (Fig. S2). Thus DIF contributes to recovery or rebound sleep following sleep loss.

**Figure 3.**
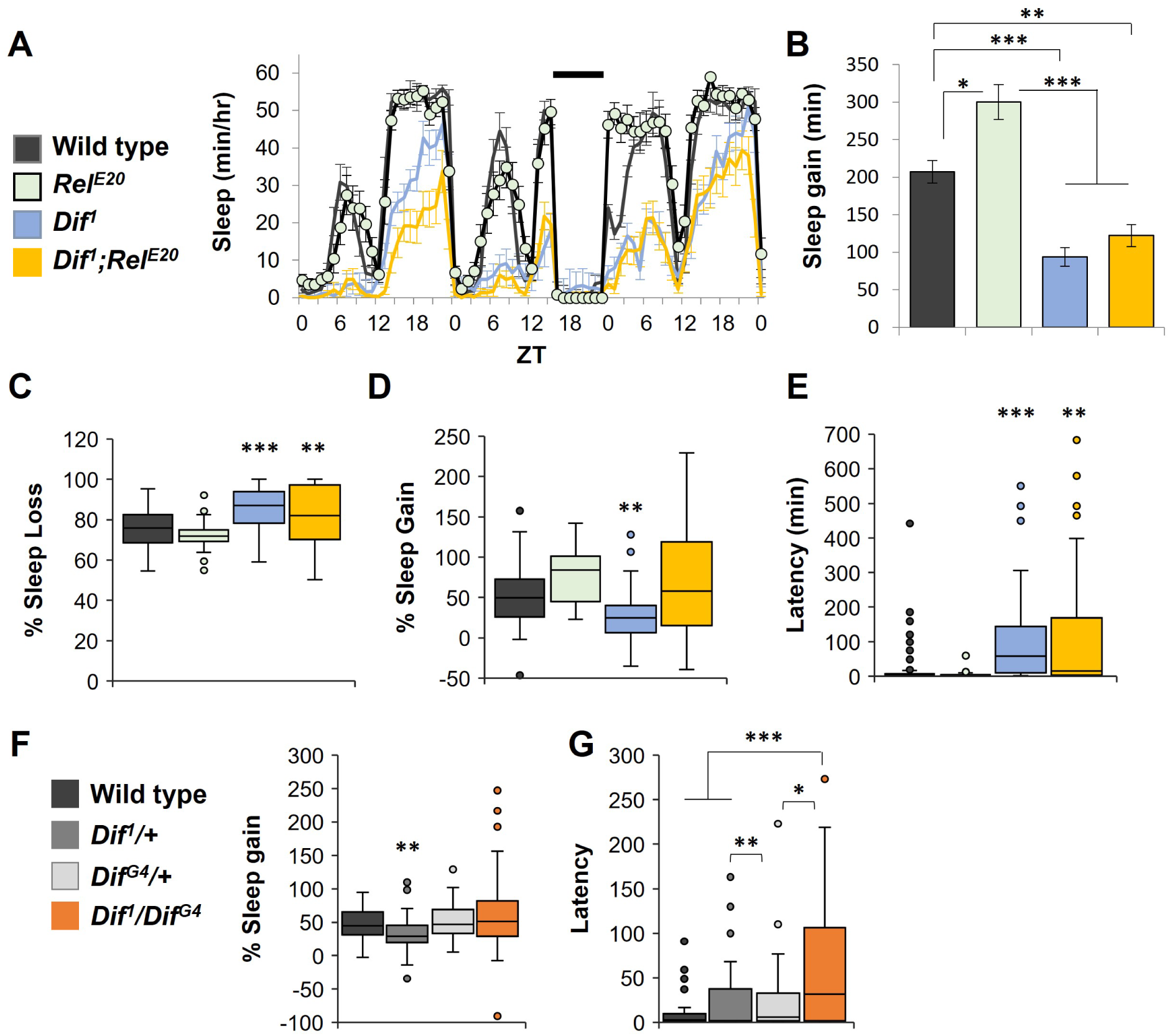
*Dif* mutants reduce recovery sleep in response to deprivation. **A**. Representative experiment showing sleep per hour (mean ± SEM; n=15-16 flies each group; ZT = zeitgeber time). Legend key applies to A-E. **B**. Sleep gain relative to corresponding baseline values during the 12h period following deprivation. *** = p<0.0001, * = p<0.01, Tukey’s post-hoc); Box and whisker plots are shown for (**C**) Percent sleep loss; (**D**) percent sleep gain; and (**E**) latency to first sleep bout following deprivation. Three outlier points in the *Dif^1^/Rel^E20^*group (468, 822, and 965% sleep gain) are not shown in (D) for better visibility. Asterisks indicate significance from CS wild type flies, *** = p<0.0001, **= p< 0.001, * = p<0.05, Kruskall-Wallis with Dunn’s post-hoc (Bonferonni-corrected). N=73 CS, 29 *Relish^E20^*, 67 *Dif^1^*, and 71 *Dif^1^*; *Relish^E20^* flies.

### Expression of *Dif* in neurons regulates sleep

We next evaluated from which tissue(s) *Dif* mediates its effects on sleep. We first tested a drug-inducible pan-neuronal knockdown of *Dif* using two RNAi lines. One strongly suppressed sleep with significant drug, genotype, and interaction effects in both male and female flies (*Dif^RNAi^ ^#30579^*, Fig S3A-D). The other RNAi line showed weaker effects when measured across a 24 hour period (*Dif^RNAi^ ^#100537^*, Fig S3E). However, when day and night were analyzed separately in the weaker RNAi line, sleep at nighttime with the pan-neuronal knockdown was significantly reduced (Fig S3F). To assess the effect of RNAi knockdown on expression of *Dif* mRNA, qPCR was used to measure *Dif* mRNA in *Dif^RNAi^ ^#30579^* and *Dif^RNAi^ ^#100537^* using a ubiquitous driver, *actin-Gal4*. *Actin-Gal4> Dif^RNAi^ ^#30579^* significantly reduced *Dif* mRNA expression relative to the parent controls (F(2,15) = 11.1, p = 0.001), whereas *actin-Gal4>Dif^RNAi^ ^#100537^* did not (F(2,13) = 0.935, p = 0.417; Figure S4). We therefore focused on the *Dif^RNAi^ ^#30579^* for further analysis.

We used the *Dif^G4^* flies to express a membrane-bound GFP, *UAS-GFP:Cd8* ^21^. Given that DIF affects baseline and rebound sleep, we considered the possibility that its expression is affected by sleep loss, and so collected brains from female flies that were either undisturbed or subjected to 6h sleep deprivation. Collections were at lights-on and brains were processed with a GFP antibody. Although modest differences in staining intensity were found between sleep deprived and non-deprived flies, we noted that the GFP signal localized to distinct neuronal groups, with strong staining in the PI; (Fig 4A). We note the limitation of using *Dif^G4^* with potential semi-dominant phenotypes. However, knockdown of *Dif* using *Dif-Gal4* was effective at significantly reducing daily sleep in both male and female flies, particularly at night (Fig4B-E and Supplementary Fig S5A). Nighttime bout duration was reduced and the time to nighttime sleep onset was increased (Fig 4B-H), similar to the *Dif^1^*hypomorphs (Fig 1).

**Figure 4.**
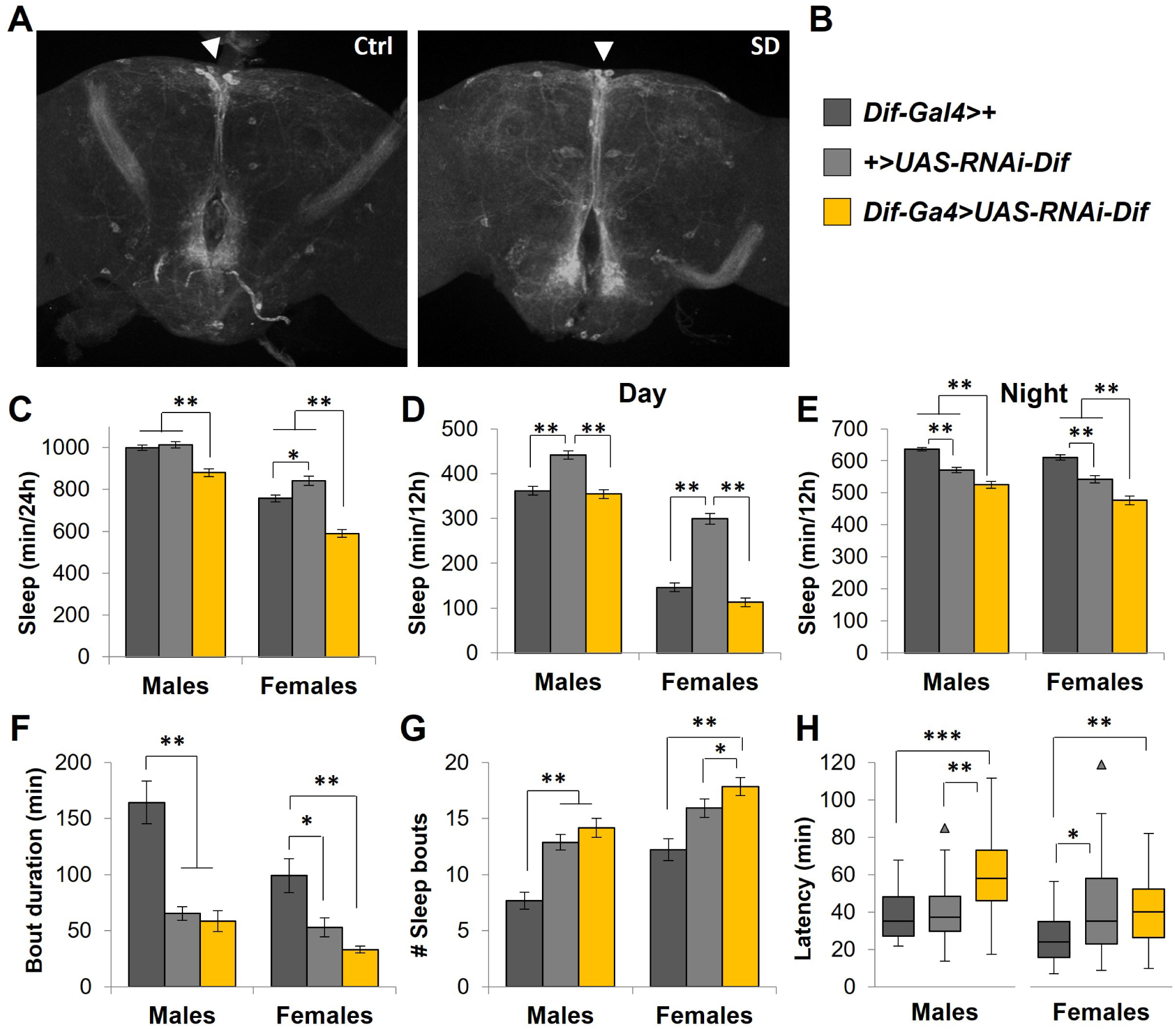
*Dif* expression pattern in brain. **A**. Confocal image of GFP driven by *Dif-Gal4* in brains from rested (“Ctrl”) or 6h sleep deprived (“SD”) female flies. Arrowhead indicates region of the pars intercerebralis. Legend key (**B**) and daily sleep profiles (**C-E**) for male and female flies of indicated genotypes; Nighttime sleep parameters, including bout duration (**F**) number of sleep bouts (**G**), and latency to first sleep bout (**H**). ***= p<0.0001; ** = p<0.001; * = p< 0.01, ANOVA with Tukey’s post-hoc in (C-G), and Kruskall-Wallis with Dunn’s post-hoc in (H); n=39-40 flies each group.

Given the high expression in the PI, we tested whether knockdown of *Dif* in these cells was sufficient to alter sleep. RNAi knockdown using *Drosophila* insulin like peptide 2 (Dilp2)-Gal4, which is known to drive expression in some PI cells ^35^, was indeed effective at significantly reducing daily sleep (Fig 5A-D and Fig S5B), particularly in females, but had no effect on recovery sleep in response to 8h deprivation (Fig. 5E and F). Knockdown of *Dif* using an alternate PI driver, c767 ^36^, showed a modest but significant effect on daily sleep in females but not in males (Fig S5C).

**Figure 5.**
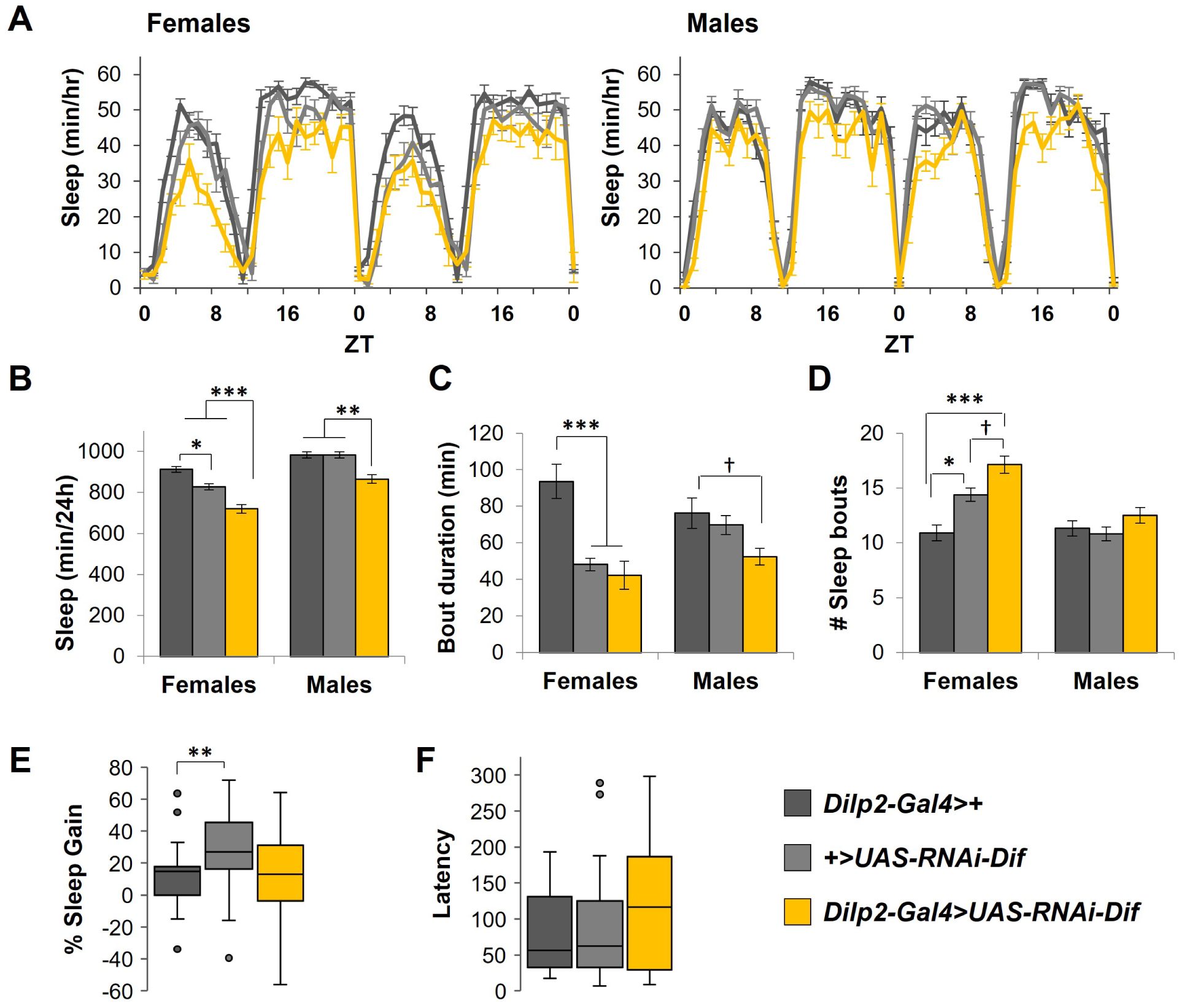
*Dif* knockdown in pars intercerebralis reduces daily sleep with no effect on recovery sleep in response to deprivation. **A**. Representative experiment showing sleep/hour in female and male flies across 2 days (n=16 flies each group; ZT = zeitgeber time). **B**. Daily sleep time in female and male flies. Nighttime sleep parameters, **C**. average bout duration, and **D**. number of sleep bouts. *** = p< 0.0001, ** = p<0.001, * = p<0.01, and † = p<0.05, one-way ANOVA with Tukey’s post-hoc; n=45-48 females, and 39-40 males each group. Box and whisker plot showing the effects of 8h sleep deprivation on daytime recovery sleep, **E**. % sleep gain, and **F**. latency (minutes) to first sleep bout after the deprivation period. Outliers are data points greater than 1.5 times the interquartile range. ** = p< 0.01, Kruskall-Wallis with Dunn’s post hoc, n=30-32 flies each group.

We therefore used a selection of alternate neuronal and non-neuronal Gal4 drivers to test a function of *Dif* in other cell groups. These Gal4 lines covered fat body (Fig S5D; Gbp3-Gal4 ^37^), glia (Fig S5E; Repo-Gal4 ^38^), mushroom bodies (Fig S5F-I ^20^, and the subesophageal region (Fig S5J-K; ppk-Gal4 ^39^ and hugin-Gal4 ^40^). Reduced expression of *Dif* in fat body decreased sleep in both sexes but did not decrease the response to sleep deprivation (Fig S6A). We also evaluated the effect of *Dif* knockdown in glial cells on the response to sleep deprivation. There was no effect on recovery sleep, nor on latency to the first sleep bout after the deprivation period (Fig S6B).

*Dif* targets several antimicrobial peptides including *drosomycin*, *bomanins*, and *metchnikowan* (*mtk*) ^41^. We reported previously that a brain expressed antimicrobial peptide, *nemuri* (*nur*), strongly promoted sleep and loss of it reduced responses to sleep deprivation ^18^. Moreover, *nur* mRNA is induced by both sleep deprivation and bacterial infection. Because baseline expression of *nur* is extremely low, we surmised that *nur-Gal4* would not be sufficient to effectively knock down *Dif* RNA. We therefore selected neuronal VT Gal4 lines ^20^ that were effective at increasing sleep when driving expression of *UAS-nur* ^18^. All four drivers tested with *Dif* RNAi were effective at significantly reducing sleep, particularly in female flies (Fig S5L-O and Fig 6A, B). We next evaluated whether knockdown of *Dif* by any of these drivers altered responses to sleep deprivation. Of these, knockdown of *Dif* using VT008975 significantly reduced recovery sleep (Fig 6C) and increased latency to the first sleep bout after deprivation (Fig 6D). While *Dif* knockdown by the other VT drivers reduced sleep, responses to sleep deprivation were variable (Fig S7A-C).

**Figure 6.**
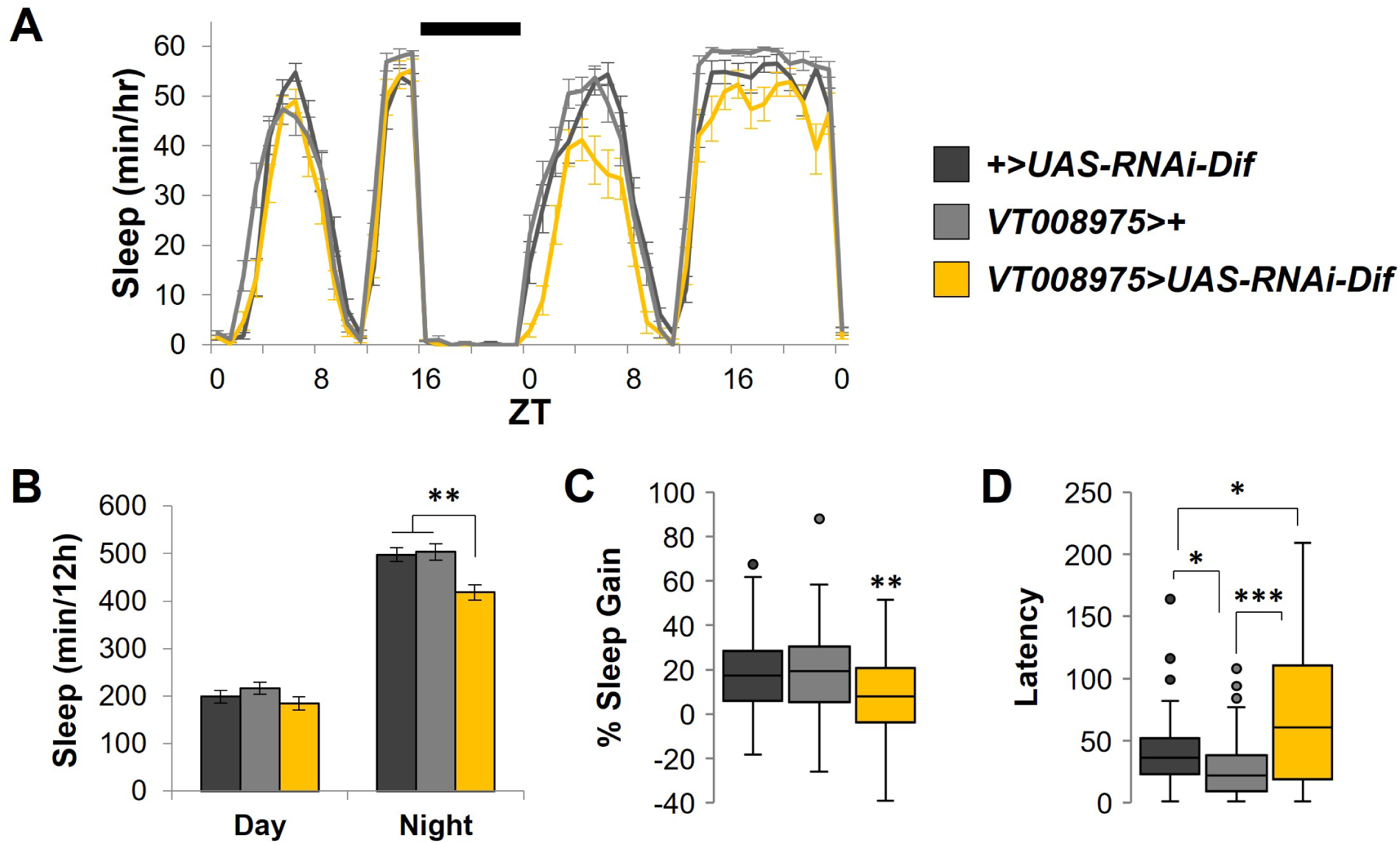
*Dif* knockdown in CNS reduces sleep and response to sleep deprivation. *VT008975*-Gal4 was used to knock down *Dif* expression. **A**. Representative experiment showing sleep per hour (mean ± SEM; n=15-16 flies each group). Horizontal bar (top) indicates the time of sleep deprivation from ZT 16-0 (ZT = zeitgeber time). Legend key applies to A-E. **B**. Daily day-night sleep values, *=p<0.01, ***=p<0.0001, one-way ANOVA with Tukey’s post-hoc, n=31-32 flies each group; **C**. Percent sleep gain measured during the 12h period after sleep deprivation; **D**. Latency to first sleep bout after deprivation; *=p<0.03, ***=p<0.0001, Kruskall-Wallis with Dunn’s post-hoc, n=22-31 flies each group for (C) and (D).

### The *nemuri* antimicrobial peptide is targeted by *Dif*

The VT Gal4 lines used for *Dif* knockdown were not only capable of increasing sleep when used to ectopically express *UAS-nur*, but also had in common projections to the vicinity of the dorsal fan shaped body (dFB; ^18^). This conclusion was based on the observation that expression of *UAS-nur* in these cells showed labeling of the NUR peptide in the dFB, indicating secretion from nearby terminals^18^. The dFB is a structure in the fly brain that is modulated by sleep pressure^42–46^. However, we note the wide distribution of labeling by each of these lines, thus the role of other cell types cannot be ruled out (Fig S7D). In any case, given that effects of *Dif* and *nur* mapped to some of the same neuronal groups, we next evaluated whether *nur* functions in an NFκB-dependent pathway. We hypothesized that if so, induction of *nur* would be abolished or reduced in mutants of NFκB, and that pan-neuronal overexpression of *nur* in a mutant background would suppress a reduction of sleep caused by loss of *Dif* in particular.

A 6-hour sleep deprivation increased expression of *nur* mRNA by 2-3 fold in both wild type flies and *Relish^E20^* mutants (Fig 7A). Normal induction of *nur* mRNA in the *Relish* mutants is consistent with the observation that these flies show recovery sleep following deprivation (Fig 3A-E). *Dif^1^*mutants, on the other hand, significantly suppressed induction of *nur* mRNA as compared to all other genotypes, including the *Dif^1^*; *Rel^E20^*double mutants which show induction intermediate to *Dif^1^* and *Relish^E20^* mutants. These findings are consistent with results reported in Fig 3D showing a significant reduction in recovery sleep in the *Dif^1^* mutants.

**Figure 7.**
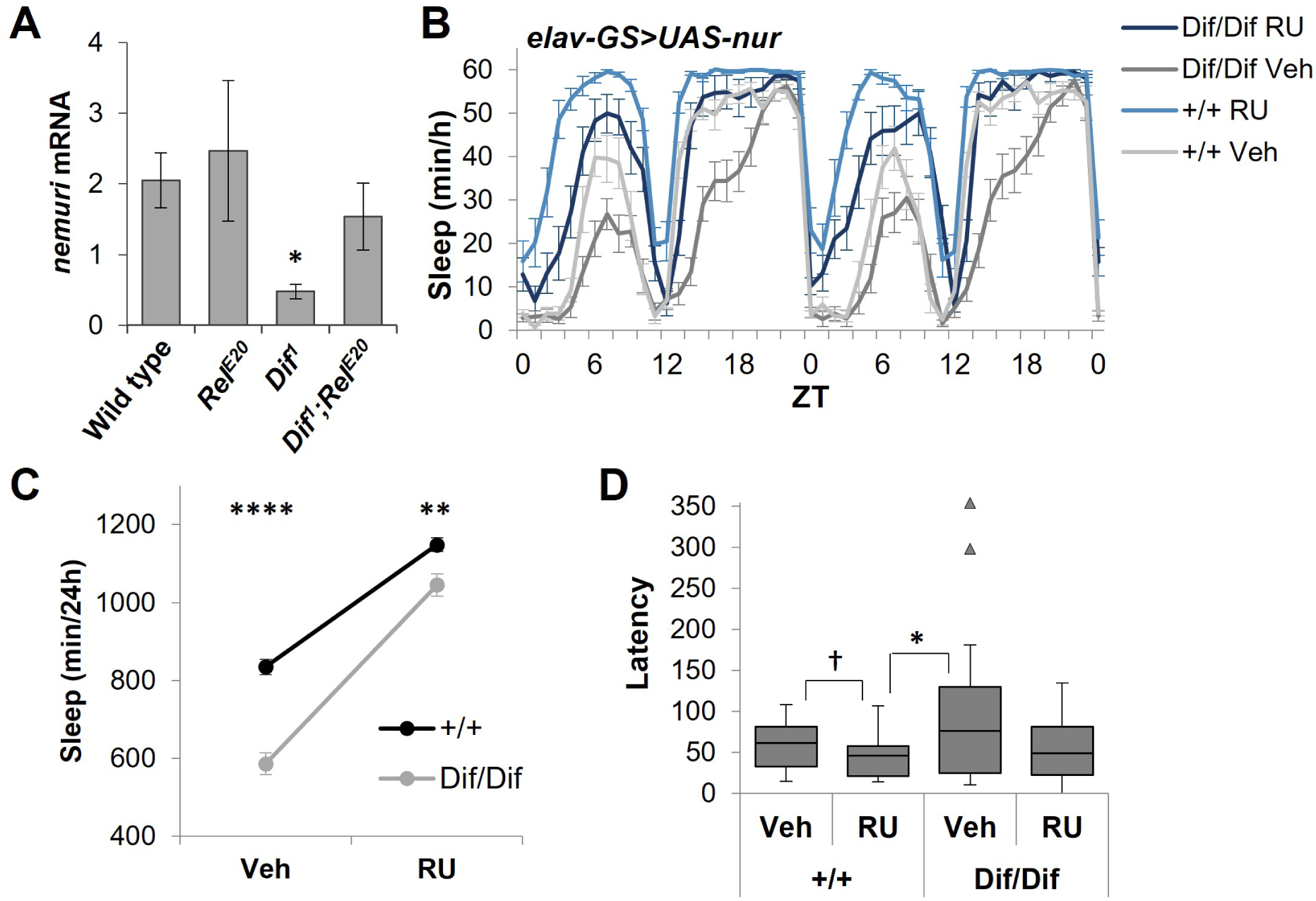
DIF targets *nemuri*. **A**. Relative *nemuri* mRNA induction by sleep deprivation in indicated genotypes, *=p<0.03, Mann-Whitney with Bonferroni correction, n= 6-7 independent experiments. **B**. Mean ± SEM sleep per hour is plotted for *elav-GS>UAS-nemuri* expressed in either wild type (*+/+*) or the *Dif^1^* mutant background (ZT = zeitgeber time). **C**. Daily 24h sleep (mean ± SEM) for each group is plotted to show interactions of drug and genotype in *elav-GS>UAS-nemuri* flies. Two-way ANOVA, **** = p<1x10^-6^, ** = p<0.006, n=37-45 flies each group. **D**. Nighttime latency to first sleep bout is shown for each group, †= p<0.05, *=p<0.005, Kruskall-Wallis with Dunn’s post-hoc.

As reported previously, conditional pan-neuronal overexpression of *nur* significantly increases sleep ^18^. Overexpression of *nur* in the same manner in the *Dif^1^* mutant background also significantly increased sleep, though not to a level equivalent to wild type controls (Fig 7B and 7C). Two-way ANOVA revealed highly significant effects of both drug and genotype (wild type versus *Dif^1^* mutants expressing *elavGS>UAS-nur*) on 24h sleep values as well as a significant interaction (F(1, 163) = 9.93, p < 0.002), where *nur* induction produced a relatively stronger increase sleep in the *Dif^1^* mutants than that in wild type controls (Fig 7C). In the wild type background, *nur* induction significantly decreased nighttime sleep latency (Fig 7D). Nighttime sleep latency trended longer in the *Dif^1^* mutants in the absence of *nur* induction, as expected. However, with drug-induced induction of *nur* in the mutants, nighttime sleep latency was indistinguishable from that in wild type flies (Fig 7D). These findings indicate that *nur* induction partially suppresses the *Dif^1^*sleep phenotype, and support the notion that *nur* functions in a sleep-promoting pathway that is downstream from *Dif*.

## Discussion

In the current study, we used *Dif* hypomorphs, which are viable in flies ^17^, and RNAi knockdown to test its role in sleep. Results showed that mutants of both sexes have reduced daily sleep and in females, suppress recovery sleep after deprivation. Knockdown of *Dif* in neurons was sufficient to reduce daily sleep and recovery from sleep deprivation. We also note that induction of *nemuri* was reduced in the *Dif* mutant background, indicating that this antimicrobial peptide is one target by which *Dif* promotes recovery sleep.

NFκB transcription factors are ubiquitously distributed across central and peripheral tissues in mammals; their involvement in physiology is wide ranging, complex, and vitally important ^5^. Given this complexity, it is unsurprising that there are very few studies that directly address a causal relationship between activation of these transcription factors and mammalian sleep. An additional complication is that genetic knockout of the mammalian homolog of DIF, p65, is lethal in mice ^47^. However, using a combination of immunohistochemical^48^, biochemical^49,50^, or pharmacological approaches ^51^, previous work has indeed indicated a role of p65 in mediating recovery sleep after acute sleep deprivation from its activity in the lateral hypothalamus ^48^ and cholinergic basal forebrain ^50,51^. Our current findings are consistent with these earlier studies and indicate a highly conserved role of the DIF NFκB in sleep homeostasis.

We previously reported a role of *Relish* in both daily ^8^ and infection-induced sleep ^9^. Interestingly, the effect of the *Relish^E20^*mutant on daily sleep was restricted to heterozygous females. Baseline sleep in the homozygous *Relish^E20^* mutant females shown in Fig 2A was indistinguishable from wild type flies, as expected. However, adding this mutant to the *Dif^1^* background further reduced sleep from *Dif^1^* mutant females but not males (Fig 1). We speculate that *Dif* may have compensated for the loss of *Relish* in females in the previous study and accounts for the strong reduction in sleep in the homozygous double mutants. Recent work has shown that heterozygous *Relish* mutants reduced mortality in response to traumatic brain injury, an effect that was influenced by genetic background and diet ^52^. Thus, *Relish* heterozygosity appears to produce unique genetic effects on sleep and other processes.

To determine the tissue(s) from which *Dif* influences sleep, we used both immunohistochemical and RNAi approaches. RNAi knockdown of *Dif* showed that it regulates sleep not only from brain, but also from fat body (Figs S3 and S5D). *Relish* also regulates both daily and infection-induced sleep from fat body ^8,9^. While the reduced sleep effect of *Dif* knockdown in fat body could be attributed to an off-target effect of the RNAi on *Relish* ^53^, this does not explain the reduced sleep that was also observed in males, since *Relish* mutants do not affect sleep in males^8^. A limitation of the current study is that sleep effects were reported mostly from one RNAi line (*Dif^RNAi^ ^#30579^*), since the other line tested (*Dif^RNAi^ ^#100537^*) was weak or ineffective on both knock-down and behavior. Activation of DIF occurs through post-translational events, such as phosphorylation, degradation of associated factors, and nuclear translocation ^10,54^. Thus it is also not surprising that we were unable to see detectable changes in GFP staining in sleep deprived flies as driven by *Dif-Gal4* (Fig 3A), as this reflects transcription of *Dif* instead of activity at the level of its protein product. Despite these caveats, significant effects of *Dif*-RNAi were indeed noted, both with the use of *Dif-Gal4* and *Dilp2-Gal4* drivers, the latter of which restricts expression to the PI.

The RNAi studies indicate that while sleep regulation by *Dif* occurs from its expression in PI, PI expression is not sufficient to alter recovery sleep from deprivation, at least not from *Dilp2*-containing PI cells. An alternate Gal4, *c767*, was even weaker, which led us to examine a role of *Dif* in alternate cell groups. Previous work had shown that increased expression of a DIF target, the antimicrobial peptide *mtk* ^41^, in glia increased daytime sleep ^55^. We were unable to detect significant effects of *Dif* RNAi in glia on daily sleep nor on recovery sleep after deprivation, suggesting ectopic effects of *mtk* reported by Dissel et al^55^. The previous study also showed that recovery from sleep deprivation is not affected by *mtk* overexpression, but its effect on sleep instead is related to short term memory formation ^55^. The effect of *mtk* knockout was not explored. Furthermore, since baseline antimicrobial peptide expression is generally low, *Dif* knockdown may also not be expected to produce a strong effect.

A more recent study showed that astrocytic glia release a Toll receptor ligand, *spatzle*, in response to sleep deprivation. This cytokine targets ellipsoid body R5 neurons to mediate responses to sleep need ^56^. Notably, the ellipsoid body (EB) is part of a sleep-regulating circuit that includes the dFB. We found that *Dif* knockdown was effective at reducing sleep from VT-Gal4 lines that were also capable of promoting sleep with overexpression of *nur*, possibly through projections to the dFB ^18^. Interestingly, our earlier study showed that putative targets of *nur* in the dFB were unique from those labeled by the widely used 23E10-Gal4 driver^42–44^. The effects of 23E10 cells have since been shown to instead be from the ventral nerve chord^57,58^. Although only one of the four driver lines altered the response to sleep deprivation when used to knock down *Dif*, we note that expression driven by the VT00875 line is more widespread than the others (Fig. S7D). Projections of this line to the central complex (which include both dFB and EB) remain to be characterized, but a function of *Dif* in sleep homeostasis from alternate neuronal groups cannot be ruled out.

Mutants of *nur* show reduced sleep in response to sleep deprivation and infection^18^. With sleep deprivation in particular, *nur* mutant flies showed a delay in recovery sleep, similar to the increased sleep latency phenotype observed in *Dif* mutants after sleep deprivation. The hypothesis that a *Dif-nur* pathway underlies recovery sleep is also supported by the results that 1) *Dif* is required for induction of *nur* mRNA with sleep deprivation, and 2) that overexpression of *nur* suppresses the *Dif^1^* sleep phenotype. However, it is expected that multiple targets of *Dif* contribute to its regulation of sleep across multiple contexts (such as learning) and more nuanced studies are needed to evaluate specific circuitry involved. In conclusion, *Dif* regulates sleep from multiple neuronal and non-neuronal cell types in a manner that is likely context dependent. The current study investigated a role of *Dif* in sleep homeostasis and identified *nur* as a target peptide.

## Acknowledgements

The authors are grateful to Tzu-Hsing Kuo and Adriana Prada for assistance with earlier stages of this project, Zhifeng Yue and Kiet Liu for technical assistance. Supported by NSF IOS #1025627, NIH #R01GM123783 and #R01NS124698 to J.A.W., A.S. is an investigator of the Howard Hughes Medical Institute.

## Disclosure Statement

Financial/non-financial disclosure: None

## Data Availability

All data are incorporated into the article and its supplementary material. Unprocessed raw data and accompanying analysis software (Insomniac3) are available upon request.

## List of Abbreviations

dFB: dorsal Fan shaped body
Dif: Dorsal immunity factor
Dilp: Drosophila insulin-like peptide
Mtk: metchnikowan
NFκB: nuclear factor binding the κ light chain in B-cells
*nur*: *nemuri*
PI: pars intercerebralis
*Rel^E20^*: *Relish^E20^*
UAS: Upstream activation sequence

## Notes

### Competing Interest Statement

The authors have declared no competing interest.

