## Supplementary Figures for "The NFκB *Dif* is required for behavioral and molecular correlates of sleep homeostasis in *Drosophila*"

### Supplementary Materials

### Supplementary Figures

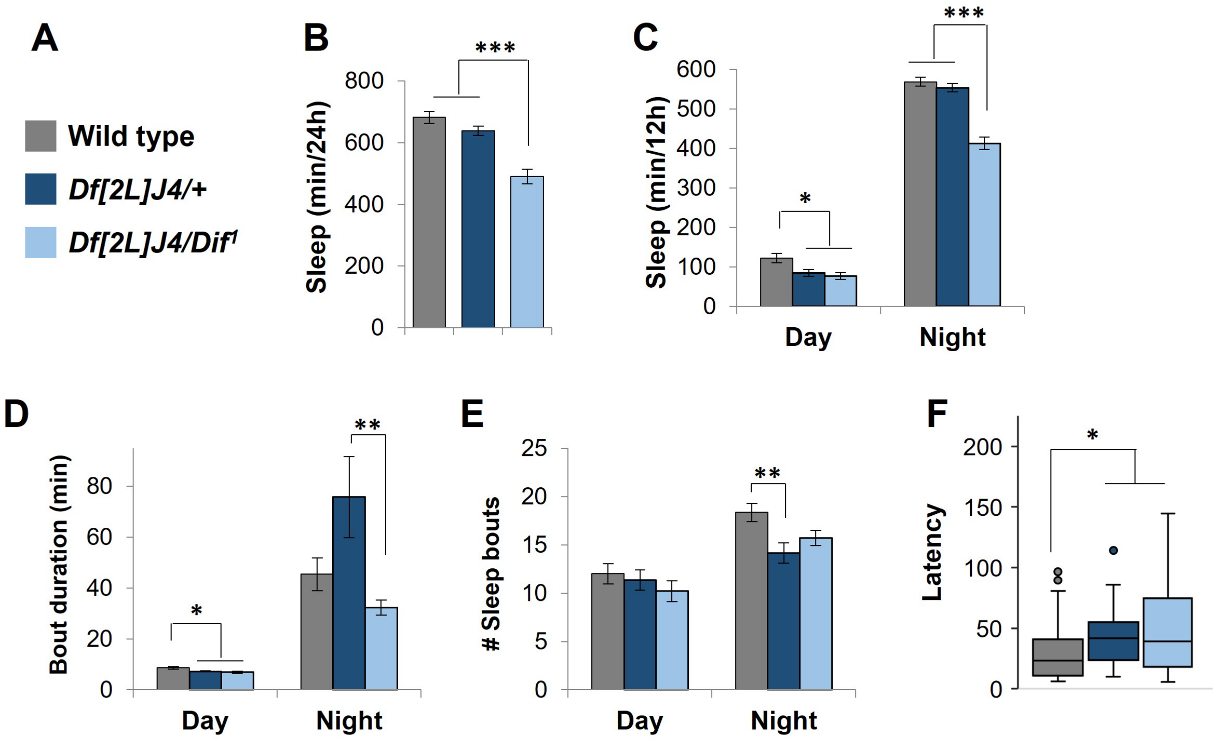


**Figure S1**. A chromosomal deficiency, *Df(2L)J4*, fails to complement the effects of the *Dif*$^{1}$ allele on daily sleep. **A**. Legend key for B-F. **B**. Mean ± SEM sleep in minutes per 24h; **C**. Mean ± SEM day and night sleep per 12h; **D**. Mean ± SEM sleep bout length and **E**. number of sleep bouts. ***= p<0.0001, **= p<0.01, *= p<0.05, one-way ANOVA with Tukey's post-hoc, n=46-58 female flies **F**. Box and whisker plot showing nightime sleep latency; one outlier at 463 min in the *Df(2L)J4/Dif*$^{1}$ group not shown, *=p<0.05, Kruskall Wallis with Dunn's post-hoc, Bonferroni corrected, n-46-58 flies.

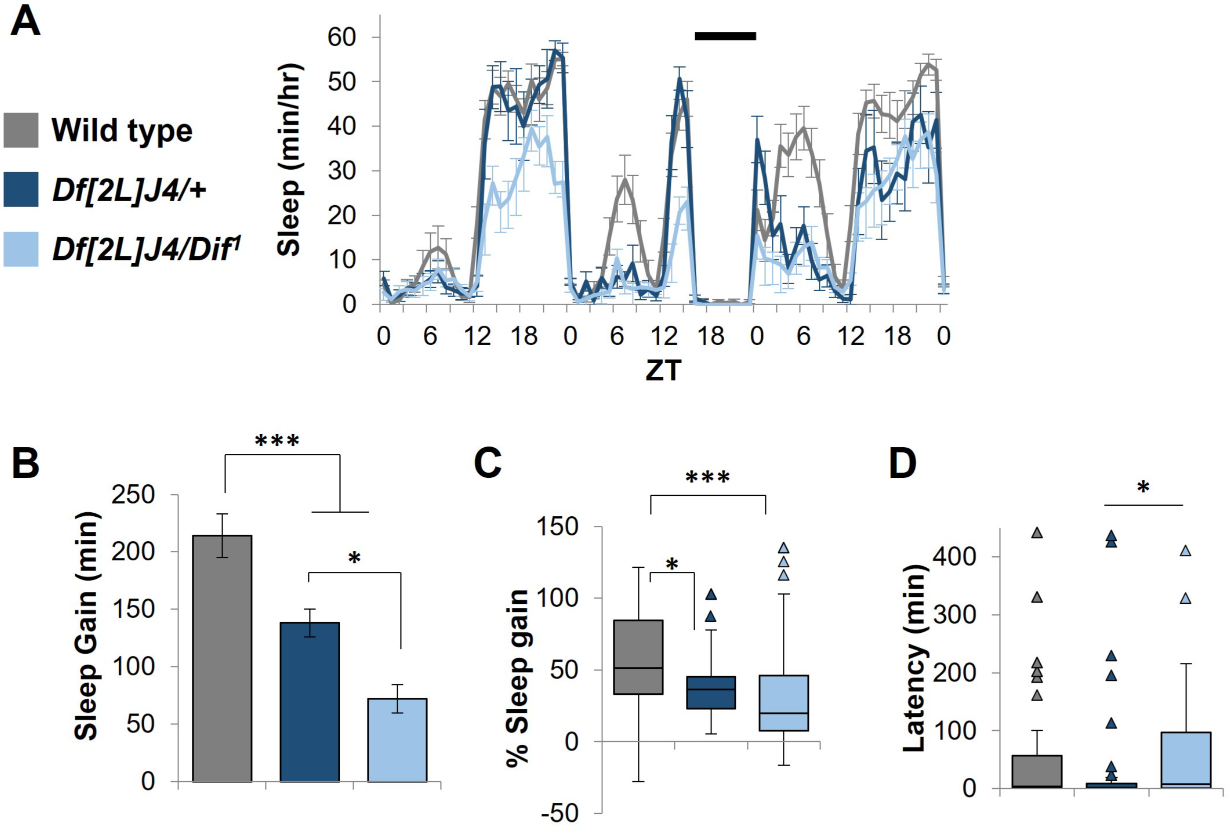


**Figure S2**. A chromosomal deficiency, *Df(2L)J4*, fails to complement the effects of the $Dif^1$ allele on recovery sleep. **A**. Representative experiment showing sleep (mean ± SEM; n=16 flies each group) per hour versus time of day (ZT) plotted for wild type, *Df(2L)J4/+,* and $Df(2L)J4/Dif^1$ flies. Legend key applies to A-D. **B**. Mean ± SEM net sleep gain in minutes during the 12h daytime recovery period after 8h sleep deprivation. ***=p<0.0001, *=p<0.01, one-way ANOVA with Tukey's post-hoc, n=30-42 flies each group. **C**. Box and whisker for percent sleep gain and **D**. sleep latency after the deprivation period. ***=p<0.0001, *=p<0.05 for both (C) and (D), Kruskall-Wallis with Dunn's post-hoc, Bonferroni corrected.

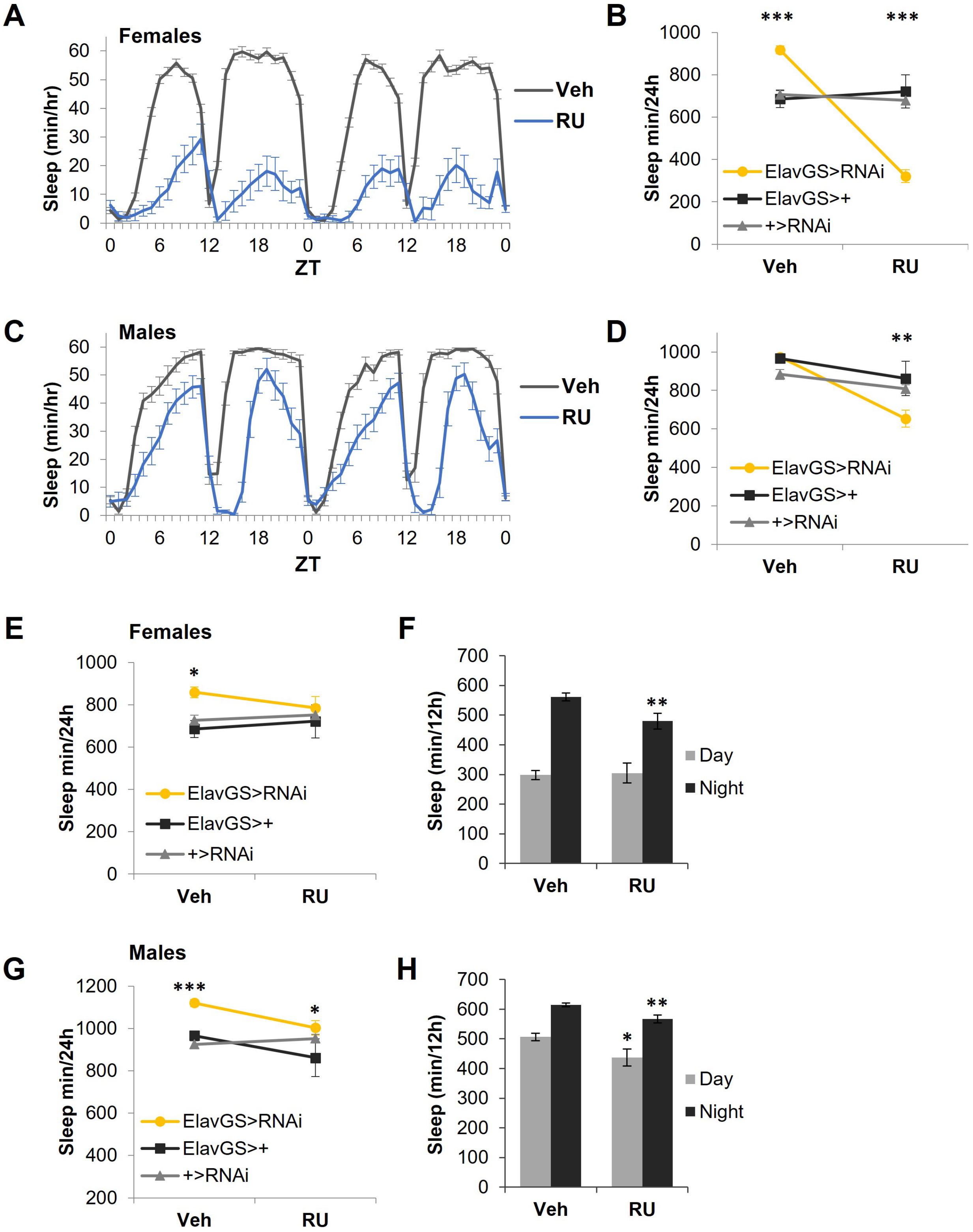
A
Females
Sleep (min/hr)
ZT
Veh
RU
B
Sleep min/24h
ElavGS>RNAi
ElavGS>+
+>RNAi
Veh
RU
***
***
C
Males
Sleep (min/hr)
ZT
Veh
RU
D
Sleep min/24h
ElavGS>RNAi
ElavGS>+
+>RNAi
**
E
Females
Sleep min/24h
*
ElavGS>RNAi
ElavGS>+
+>RNAi
F
Sleep (min/12h)
Day
Night
**
G
Males
Sleep min/24h
***
*
ElavGS>RNAi
ElavGS>+
+>RNAi
H
Sleep (min/12h)
Day
Night
*
**

**Figure S3**. Panneuronal knockdown of *Dif* reduces sleep. *elav*-geneswitch (*elav*GS) was used for drug-dependent RNAi knockdown. Flies were fed RU486 (RU) or an equivalent amount of vehicle (Veh). **A**. Representative experiment showing mean ± SEM sleep per hour in *ElavGS>Dif*$^{RNAi\ \#30579}$ female flies (n=16 flies each). **B**. Daily sleep values are reported as minutes/24h to show significant interactions between drug and genotype, including each of two control parent lines (*ElavGS/+*; n=48 and 24 for Veh and RU conditions, respectively, and *+/Dif*$^{RNAi\ \#30579}$; n=45 and 48 flies for Veh and RU conditions, respectively) and the experimental *ElavGS>Dif*$^{RNAi\#30579}$ line in yellow (n=46 Veh and 47 RU flies). ***=p<0.0001, 3x2 two-way ANOVA (genotype versus drug condition). **C**. Representative experiment as in (A), in male flies (n=15 flies each). **D**. Daily sleep values in male flies, *ElavGS/+* (n=47 Veh, n=12 RU); *+/Dif*$^{RNAi\#30579}$ (n=44 Veh, n=45 RU) and *ElavGS> Dif*$^{RNAi\ \#30579}$ (n=48 Veh and n=46 RU flies). **=p<0.005, 3x2 two-way ANOVA for interactions between drug and genotype. **E.** Daily sleep values for *ElavGS/+* (same flies as in (**B**)), *+/Dif*$^{RNAi\#100537}$ (n=43 Veh, 37 RU), and *ElavGS>Dif*$^{RNAi\#100537}$ (n=47 flies each for Veh and RU groups). *=p<0.02, 3x2 two-way ANOVA (genotype versus drug condition). **F**. Sleep (min/12h) is shown for the same groups in females; **= p<0.008 compared to the nighttime value in the Veh control group, student's *t*-test. **G.** Daily sleep values per 24 h for males for *ElavGS/+* (same flies as in (D)), *+/Dif*$^{RNAi\#100537}$ (n=46 Veh, 40 RU), and *ElavGS>Dif*$^{RNAi\#100537}$ (n=48 and 46 for Veh and RU, respectively), ***=p<0.0001, *=p<0.03, 3x2 two-way ANOVA (genotype versus drug condition). **H**. Sleep (min/12h) is shown for the same groups in males (**G**); *=p<0.03, **=p<0.002, student's *t*-test comparing day and nighttime values, respectively.

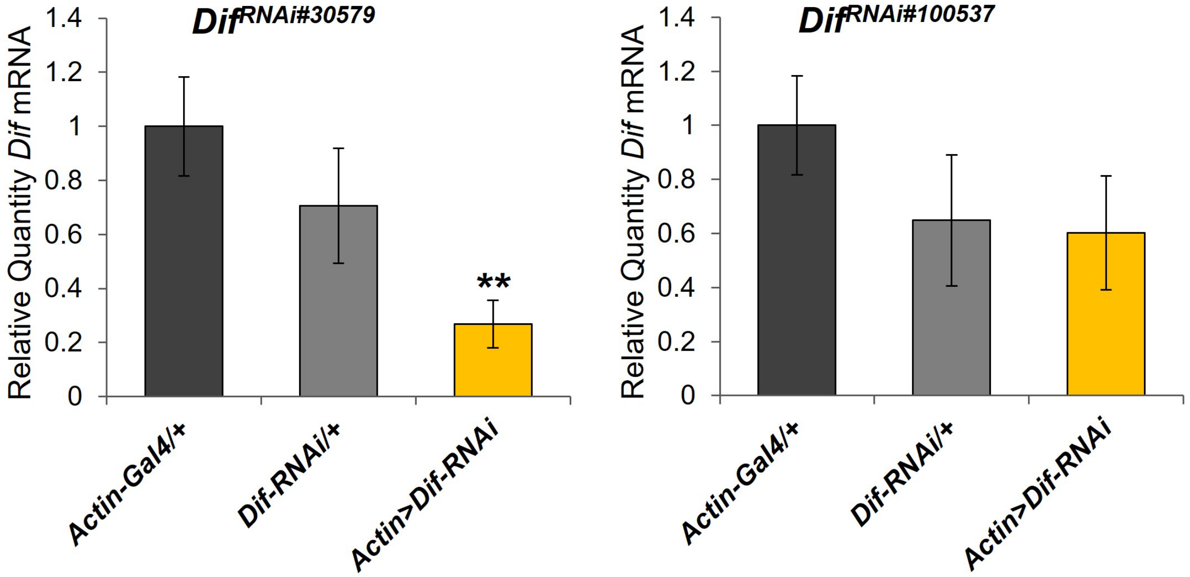


**Figure S4**. Effect of *Actin-Gal4>UAS-Dif RNAi* on expression of *Dif* mRNA. Relative quantities of *Dif* mRNA (mean ± SEM) in indicated genotypes. **=p<0.005, one-way ANOVA with Tukey's post-hoc.

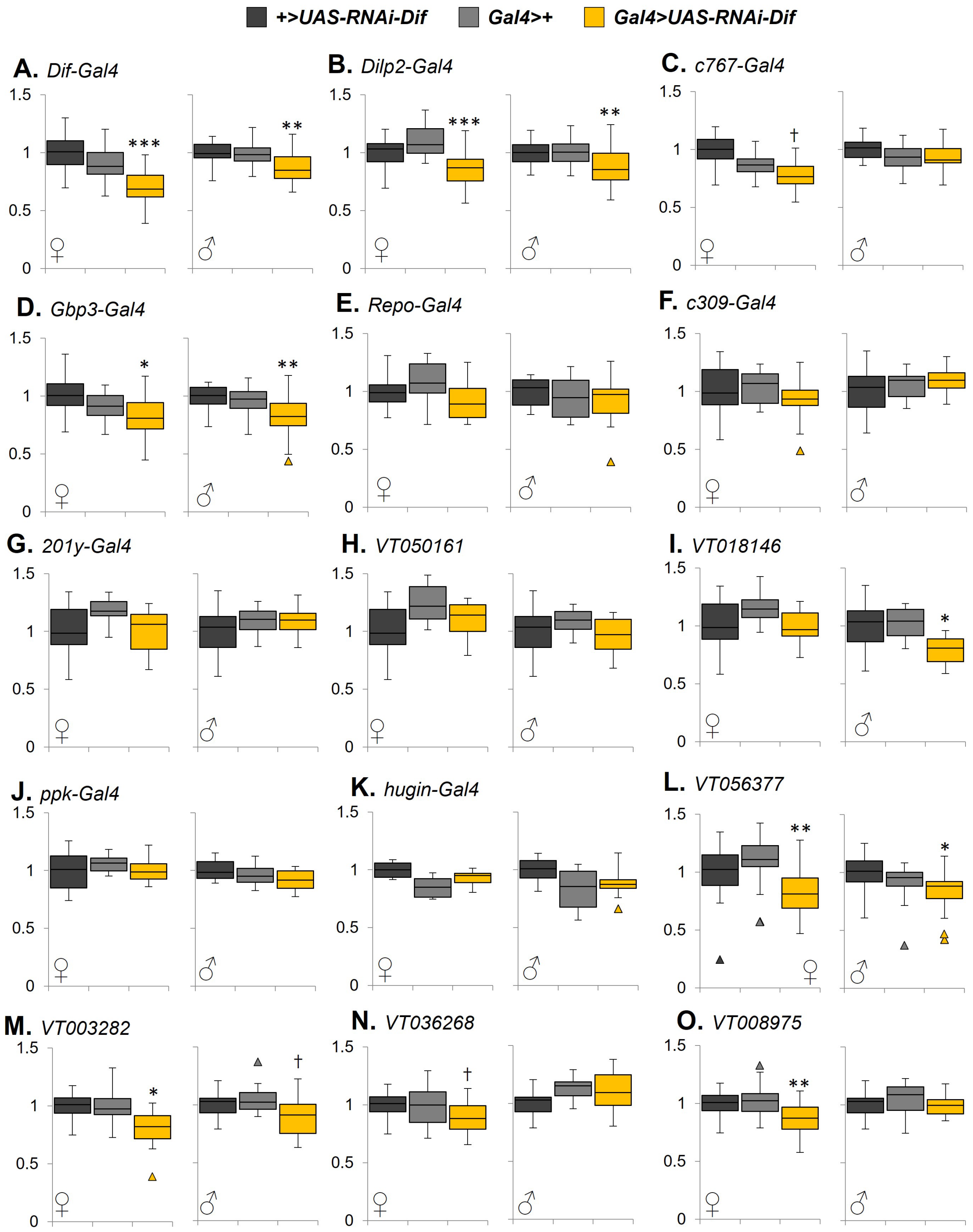
+>UAS-RNAi-Dif
Gal4>+
Gal4>UAS-RNAi-Dif
A. Dif-Gal4
B. Dilp2-Gal4
C. c767-Gal4
D. Gbp3-Gal4
E. Repo-Gal4
F. c309-Gal4
G. 201y-Gal4
H. VT050161
I. VT018146
J. ppk-Gal4
K. hugin-Gal4
L. VT056377
M. VT003282
N. VT036268
O. VT008975

**Figure S5**. Effects of *Dif* knockdown in neuronal and non-neuronal cells with *Gal4* drivers indicated in each figure panel. Daily sleep values (minutes sleep/24h) were normalized to the control parent line, *UAS-RNAi-Dif/+* and medians with interquartile intervals (box and whiskers) are shown for male and female flies in A-O. Statistical effects are indicated *only* when the knockdown line (yellow) varied significantly from *both* parent control lines; ***=p<0.0001, **=p<0.001, *=p<0.01, and †=p<0.05, Kruskall-Wallis with Dunn's post-hoc, Bonferroni corrected. N=6-48 flies per group.

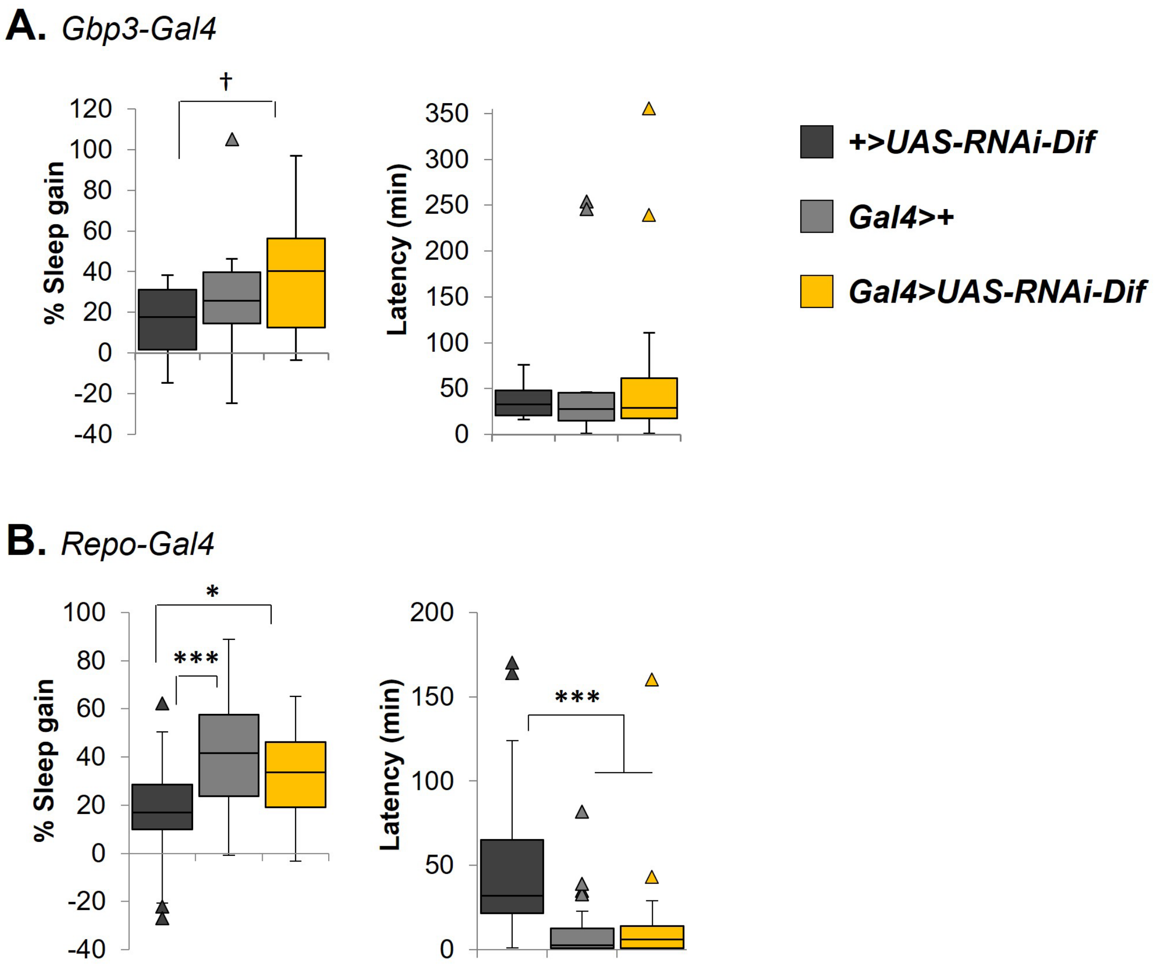


**Figure S6**. *Dif* does not affect sleep homeostasis from non-neuronal cells. Flies were sleep deprived for 8h and effects on recovery sleep are shown, including percent sleep gain (left panels) and time in minutes to first sleep bout after the deprivation (latency; right panels) for **A**. *Dif* knockdown in fat body using Gbp3-Gal4 (n=13-16 flies), and **B**. *Dif* knockdown in glia using Repo-Gal4 (n=30-32 flies each group). ***=p<0.0001, *=p<0.01, and †=p<0.05, Kruskall-Wallis with Dunn's post-hoc.

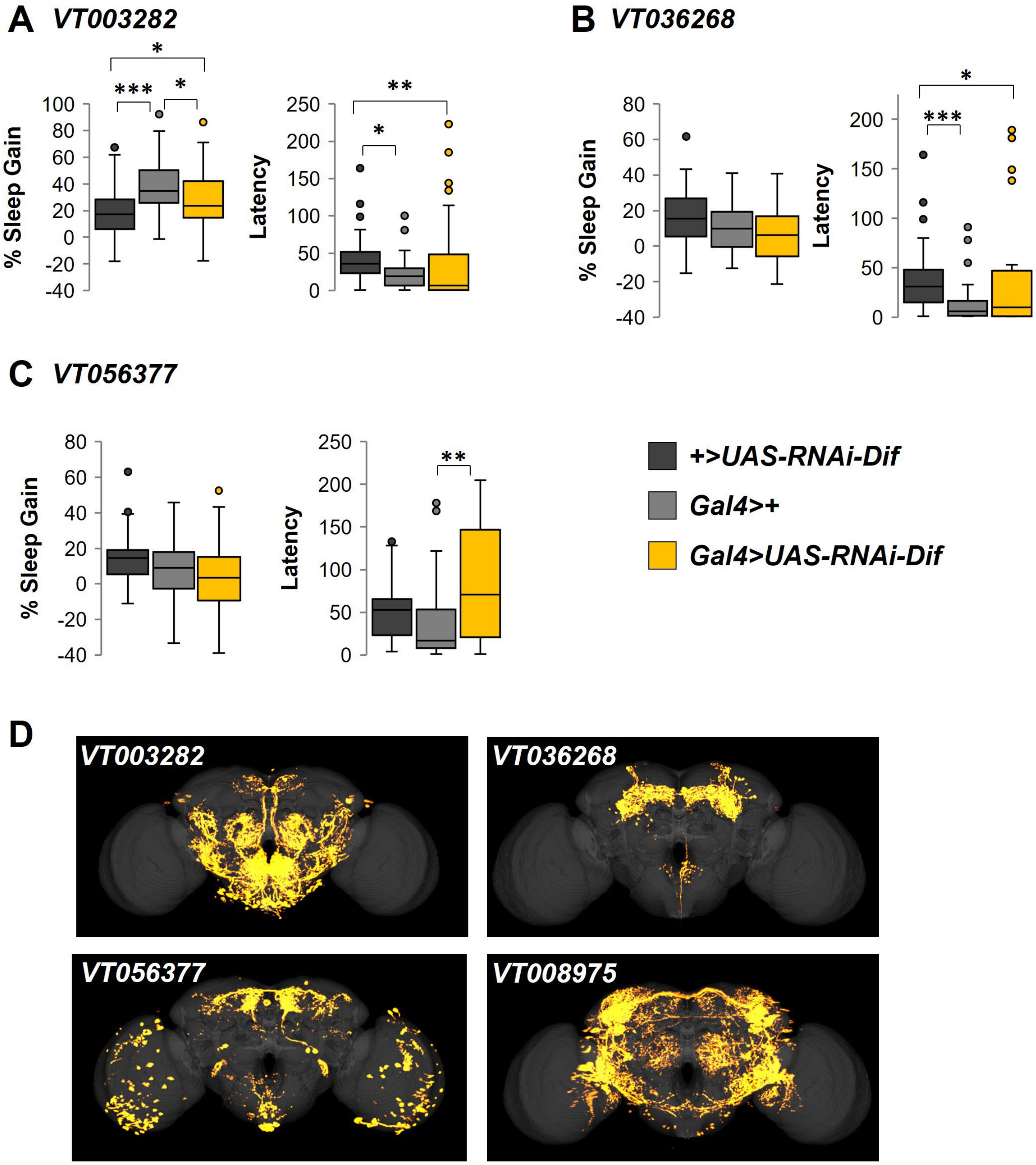


**Figure S7**. Effects of *Dif* knockdown on sleep from select Vienna Tile lines. Percent sleep gain and latency to daytime sleep onset following sleep deprivation for **A**. VT003282 (n=18-29); **B**. VT036268 (n=24-31); and **C.** VT056377 (n=32 each group). *=p<0.05, **=p<0.01, ***=p<0.001, Kruskall-Wallis with Dunn's post-hoc. **D**. Distribution of brain expression shown for indicated VT-Gal4 lines. Images originated from Tirian et al [20] and were generated from www.virtualflybrain.org [59].
